# Shadow montage and cone-beam reconstruction in 4D-STEM tomography

**DOI:** 10.1101/2025.09.06.674643

**Authors:** Shahar Seifer, Lothar Houben, Michael Elbaum

## Abstract

Diffraction images in a scanning transmission electron microscope (STEM) provide a real-space projection of the sample at sufficient probe defocus. These so-called shadow images can be acquired patch by patch in a 4D-STEM setup using a pixelated detector and assembled into a shadow montage. Due to parallel acquisition within each illuminated patch, an upscaled bright field (BF) image is rendered efficiently in time and with little additional computational demand compared to other STEM techniques. We show that in this shadow regime described by geometrical optics, the algorithm achieves the result of a tilt-corrected bright field image. Furthermore, the solution is equivalent to cone-beam reconstruction in a particular scenario of a scanning point illumination source in a plane. The contrast transfer is similar to that of conventional wide-field TEM, but like STEM the focus is insensitive to energy loss and objective lens chromatic aberration. By adjusting the overlap between shadow patch images in the diffraction plane, the shadow montage is synchronized to specific layers in the sample, rendering a 3D shadow volume from a single dataset. The method is also amenable to conventional tilt tomography, by adding a shadow montage or shadow volume to each tilt view prior to back-projection. This approach effectively circumvents the basic presumption of parallel-projection tomography that the depth of field must be greater than the specimen thickness.

## Introduction

Investigation of beam-sensitive specimens by electron microscopy depends on optimization of data acquisition modalities. Such “low dose” imaging was pioneered for studies of biological macromolecules embedded in vitreous ice. Life science cryogenic transmission electron microscopy (cryoEM) established a highly successful workflow for high-throughput data collection, which, combined with extensive image processing and averaging, routinely yields structures at atomic or near-atomic resolution. Standard practice employs wide-field parallel illumination with a large defocus, requiring at least an approximate aberration correction in software in order to achieve a reliable representation of material densities. While illumination fluences of only tens of electrons per square Angstrom (*e*^−^/Å^2^) are employed, the final structure normally includes image contributions from thousands to millions of individual particles. A fundamental presumption rests in the weak phase object (WPO) approximation, i.e., that the specimen is essentially transparent. A current challenge is to extend the workflow for cryo-EM to tomography of thicker specimens, for which this presumption becomes problematic. Moreover, electron energy loss in the specimen requires the use of an energy filter to avoid a haze that originates in the objective lens chromatic aberration. In parallel, a growing area in materials science addresses beam-sensitive specimens. These developments motivate exploration of new image acquisition and analysis workflows, many of which are based on scanned probe, i.e., scanning transmission EM (STEM) rather than wide field modalities.

A basic STEM configuration illuminates a thin sample with a converging probe, as shown in Figure 1a (Williams and Carter, 2009). Owing to the condenser lens (lens 1) and aperture, a smooth probe is formed with a radius determined by wave diffraction and lens aberrations. For a probe focused on the sample, the diverging illumination that is projected onto the detector plane is essentially uniform. The key parameter is angular aperture, i.e., the semi-convergence angle α that defines the illumination cone. The intersection of the illumination with the detector plane defines an on-axis bright-field disc. With a specimen in place, scattering away from this disc will lower the integrated intensity within. Conversely, the area surrounding this disc defines a dark-field region that is illuminated only by scattering from the specimen. Thus, detectors in the form of a disc on axis and a centered annulus acquire bright and annular dark field signals (BF & ADF), respectively. The numerous acronyms for STEM refer to specific detector configurations that define a region of the diffraction plane over which the transmitted signal is integrated.

**Figure 1.**
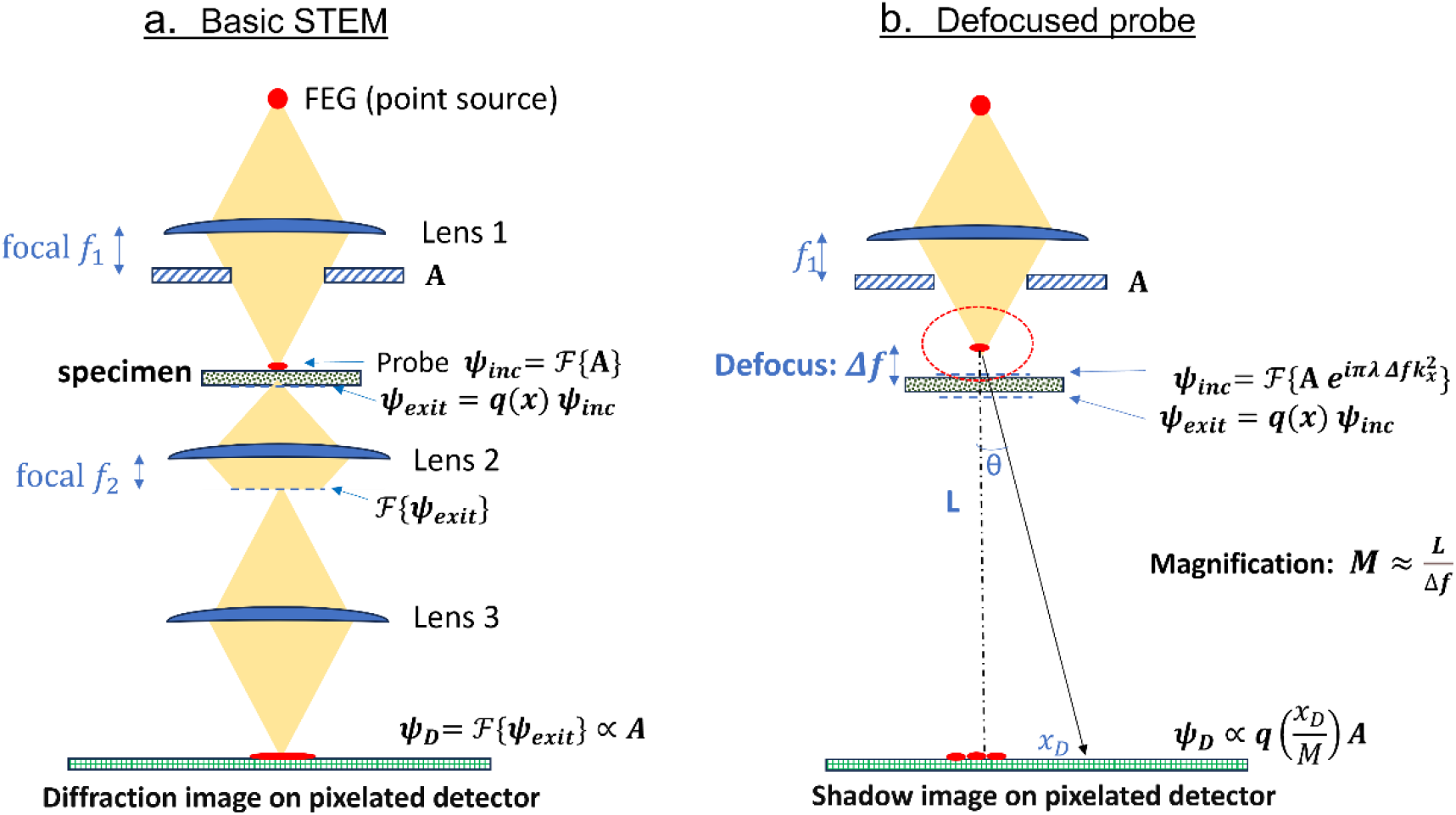
Simplified scheme of the basic STEM and the defocused probe configuration. **(a)** The condenser lens (Lens 1) focuses the probe onto a thin sample. Lenses 2 and 3 are equivalent to a far field diffraction of the exit wave, which appears as a “flat” BF disc. **(b)** Defocused probe introduces phase aberration for which the wavefunction at the detector contains real space structural details of the sample.

We have shown previously that very thick biological specimens, up to one micron or more, may be studied by scanning transmission EM (STEM) (Elbaum, 2018; Rez et al., 2025; Wolf et al., 2014). The classical STEM configurations using single-element detectors do not exploit the wave coherence, in the sense that momentum information is lost. Recent innovations have introduced methods to generate position-momentum contrast, ranging from differential phase contrast (DPC), and derivatives thereof, to full iterative ptychography (Ding et al., 2022; Küçükoğlu et al., 2024; Lazić et al., 2022; Lazić and Bosch, 2017; Maiden and Rodenburg, 2009; Seifer et al., 2021). While DPC may be implemented using simple quadrant detectors, more sophisticated methods require pixelated detectors that can record a substantial part of the diffraction plane per pixel in real space. This is the essence of 4D STEM (Ophus, 2019). The approach gathers momentum as new fast detectors become available. In parallel, there is important activity to evaluate the most suitable analytical tools to handle the very large but sparse datasets generated under low-dose imaging conditions. A notable recent advance is that of tilt-corrected bright field (tcBF), or parallax STEM imaging, which employs a defocused but still highly coherent probe to acquire from an area substantially larger than the diffraction-limited probe that limits resolution (Küçükoğlu et al., 2024; Yu et al., 2024). The defocus aberration is corrected by imposing image shifts on the images generated from individual sampled pixels of the diffraction plane; the multi-pixel collection over the defocused patch enables acquisition of a large field of view in reasonable time. The image shift analysis bears a strong resemblance to parallax-filtered integrated DPC using a quadrant detector (Seifer et al., 2021, 2024b). In this work we present an alternative to tcBF based on a montage of shadow images that appear in the diffraction plane, including application to tomography. Relative to iterative ptychography the image processing is straightforward and direct. In common with those methods, the approach effectively circumvents the conventional trade-off between scan resolution and field of view.

Formally, the wavefunction at the detector plane is a 2D Fourier transform of the exit wave at the bottom of the sample, therefore equivalent to a far field diffraction. In practice, the diffraction pattern produced at the objective back focal plane is magnified by the microscope projection optics to mimic a free-space transmission over a distance *L* that is called the camera length. *L* relates the radius on the detector plane to the angle with respect to the optic axis, or equivalently, the momentum vector *k*. Thus, the illumination cone describes a cutoff *k<k’*, which also defines the bright field detector radius. For an image scanned in focus, the optical resolution is determined by the diffraction-limited probe radius *λ*/*α*, while the depth of field scales as *λ*/*α*^2^. (Spherical and higher order aberrations are ignored here for simplicity.) The depth of field is particularly relevant for tomography, which is traditionally based on a parallel projection presumption.

A defocused probe as shown in Figure 1b introduces more structural information in the detector wavefunction. The defocused probe obviously covers a larger area of the specimen. For a thick specimen and large convergence, the illuminated radius will be a function of depth. Some ptychography methods such as ePIE (Maiden and Rodenburg, 2009) reconstruct the exit wave and the probe function from their convoluted power spectra on the detector, assisted by precise overlap of translated probe in real space. For specific optical specimens, the ptychography approach has demonstrated deep sub-Angstrom resolution limited by lattice vibration rather than physical optics (Chen et al., 2021). However, ptychography relies on wave coherence that may have limited applicability to thick amorphous samples, and iterative methods of phase retrieval may not converge robustly under the sparse exposure conditions relevant to biological cryo-EM. In that realm, an encouraging recent work demonstrated sub-nanometer resolution for protein specimens by combination of iterative ptychography with single particle analysis (Küçükoğlu et al., 2024).

In a prescient paper, Cowley (Cowley, 1979) showed that for a defocused probe the far-field diffraction patterns are essentially shadow images of the sample, magnified by the ratio of camera length to defocus (see example in Supplementary Video 1). Owing to the phase imposed on the incident wave, a shadow of the sample is formed within the BF illumination disc. Precisely such shadow images under a defocused probe are collected as raw data in a 4D-STEM measurement. We show here that a large 2D montage can be constructed simply from the scanned probe samplings, and moreover that depth information can be recovered due to the strong dependence of magnification on defocus. The shadow montage method is further applied to tilt tomography to generate a 3D reconstruction with a large field of view and lateral resolution that is not constrained by the parallel projection presumption. We expect that this direction has significant potential to become a standard imaging tool for electron microscopy of beam-sensitive specimens, and in particular biological cryo-EM.

Assembly of the shadow montage from individual shadow patch images is demonstrated in Figure 2. The intensity value in each pixel is normalized by the number of sourced probe positions, in order to account for overlap of the illuminated patches. Several shadow images are produced simultaneously with different synchronization steps, namely, the distance (in pixels) between adjacent patches of camera images. This synchronization step depends on the step size between probe centers in the scan, and on the magnification of the shadow image projected onto the diffraction plane. The next stage is to down-sample the images to a size consistent with the actual optical resolution limit for the given de Broglie wavelength *λ* and semi-convergence angle α. Resolution may also be limited by Poisson noise at low dose (Egerton, 2024), as well as the characteristics of a display or analysis tool. In the final stage we remove inversions (sign-flips) in the contrast by working on the reciprocal representation of the image, similarly to standard contrast transfer function (CTF) processing for TEM (Wan et al., 2012). CTF correction is essentially a deconvolution process aiming to balance the spectrum of the signal response.

**Figure 2.**
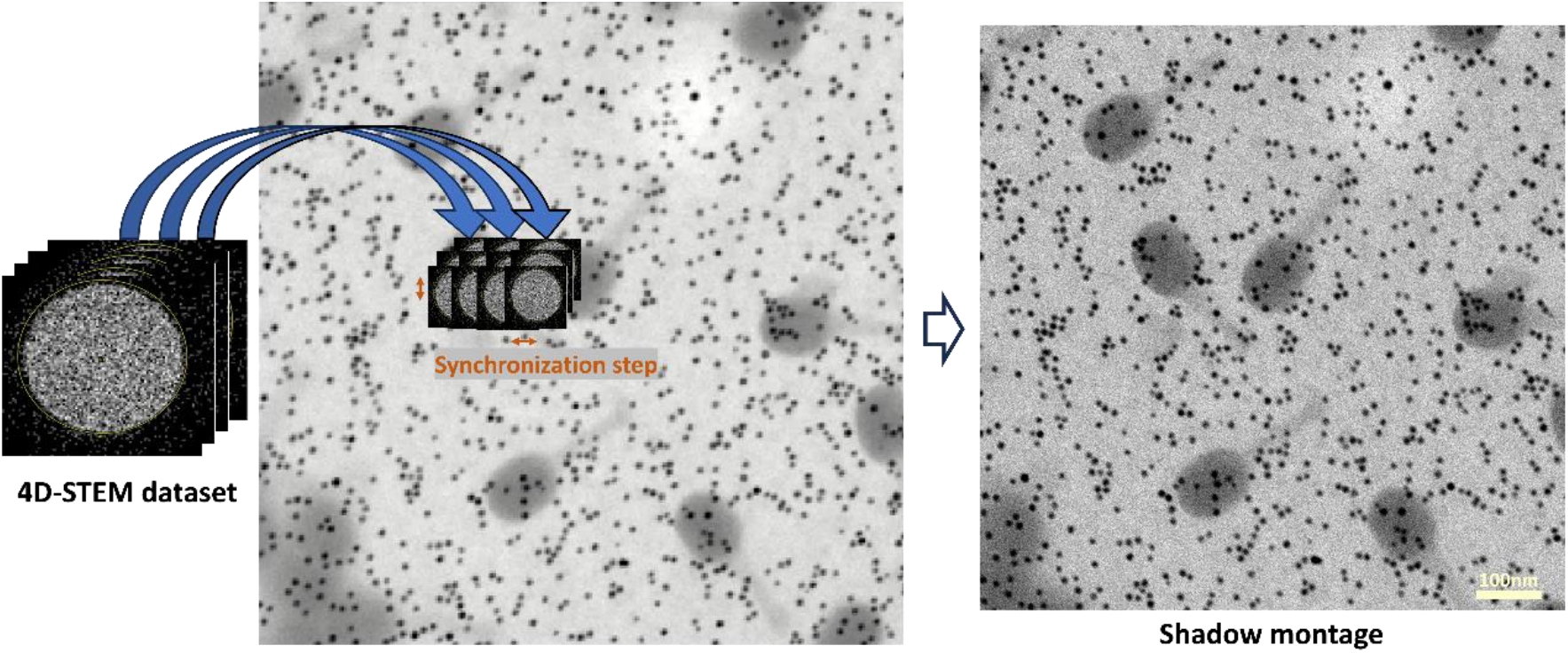
Generation of a shadow montage from overlapped camera images at a certain synchronization step. The basic BF-STEM image is shown on the left for demonstration. The sample consists of gold nanoparticles and T4 bacteriophages.

In spite of the defocus, the shadow montage potentially offers much higher resolution than a basic BF-STEM image (compare the images in Figure 2). This is achieved by inverting the parallax effect, meaning that each pixel in the detector plane is reassigned to a neighboring probe location according to the aberration function. Tilt-corrected bright field (Yu et al., 2024, 2021) is an ultimate extension of this approach: it generates up-sampled images based on reassignment of the diffraction camera pixels to interpolated positions of the scanned probe in real space. The tcBF algorithm also correlates between the scan images generated by individual camera pixels in order to calibrate the aberration function in each case. In that sense it is in principle extensible to aberration of any order. In the special case that a defocus aberration is dominant, the tilt correction (equivalently, the parallax shift) is simply the product of the defocus with the coordinate of the camera pixel relative to the illumination center. Such a linear correction is equivalent to a condition of uniform magnification in the shadow image. This simplifies the problem considerably and allows generation of the tcBF from all the camera pixels at once by shifting each shadow image to its appropriate location in a large up-sampled montage (see example in Supplementary Figure S1). This procedure provides an image equivalent to the tcBF in a way that is both intuitive to understand and fast to implement.

Shadow montage and CTF correction follow a three-step strategy in optimal signal recovery: synchronization, summation, and deconvolution (SSD). Synchronization is most intuitive for stitching of the shadow images into a large montage; the overlap between small illuminated patches achieves the summation. SSD is also a useful concept in tomography. We showed previously that an optimal 3D reconstruction is achieved by back-projection followed by deconvolution (Kirchweger et al., 2023; Seifer et al., 2024b; Waugh et al., 2020). The back-projection is essentially a process of rescaling and shifting of a tilt series followed by summation. The process is demonstrated in details in Supplementary Video 2 for a lignin sample (Palakurthy et al., 2025). A linear transformation synchronizes the apparent drift of objects in the projection of a certain layer as a function of tilt angle along the sinogram. After such transformation the chosen plane becomes stationary, and the contribution of the different projections can be summed self-consistently to generate contrast in that particular layer. Repeating the process for all layers fills the back-projected volume. We then further deconvolve the result with a characteristic kernel to suppress artifacts that result from the limitation of discrete projection angles. These appear as rays or spines “fanning out” from points of high contrast. A sampling theory approach offers certain choices of deconvolution kernel for optimal reconstruction (Seifer, 2023). The SSD concept is also apparent in cone-beam reconstruction techniques common in medical CT imaging (Bronnikov, 2000), which we adapt additionally for cryo-tomography.

### Theory

Following Cowley (Cowley, 1979), we consider a thin sample illuminated by a STEM probe at a certain defocus Δ*f*. The incident wavefunction entering the sample is distorted by an aberration 𝒳, a function of the incident *k* vector components. 𝒳 represents minimally the defocus, converging or diverging, as well as higher-order contributions. A position dependent function *q(x,y)* describes the complex specimen transmission function that includes imaginary and real parts, i.e., phase and amplitude modulations. The transmission function effectively introduces refraction of the rays at the sample. The propagation is described in two steps (see Figure 1b): from a condenser aperture (CA) to the sample with aberration 𝒳, and from the sample to the diffraction plane. The latter is often drawn and modeled as a free space propagation; in fact, it involves a further magnification in the projection system, whose possible additional aberrations we neglect. Thus, the wavefunction at the detector plane *ψ*_*D*_ is found as follows

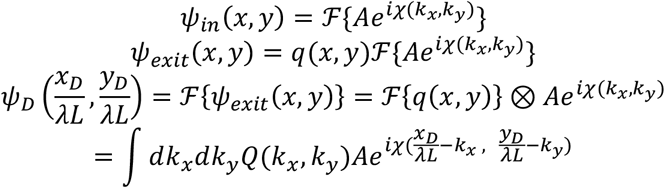

Examining a point (*x*_*D*_, 0) at the diffraction plane (without loss of generality) and expanding the aberration function term to first order in *k*_*x*_

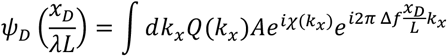

This expression is interpreted as a Fourier transform, provided that the aperture in *k*_*x*_ allows integration of at least one cycle in the last exponent, namely 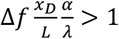. It means that for an artifact-free shadow image the defocus Δ*f* should be larger than the depth of field *λ*/*α*^2^ (see table 1). At such condition

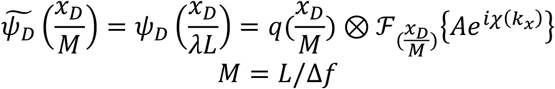

**Table 1.** Recommended lower bounds on defocus for various convergence angles (200 kV, C_s_=2 mm, Δx_S_= 2 nm).

| criterion | $\alpha=0.8$ mrad | $\alpha=4$ mrad | $\alpha=9$ mrad |
| --- | --- | --- | --- |
| $\lambda/\alpha^2$ | 3900 nm | 150 nm | 31 nm |
| $C_s \alpha^2 / 0.1$ | 13 nm | 320 nm | 1620 nm |
| $\Delta x_S / 2\alpha$ | 1250 nm | 250 nm | 110 nm |

The right-hand side can be read with *x*_*D*_ replaced by the real space coordinate *x* = *x*_*D*_/*M*. Namely, due to a certain defocus Δ*f* and a camera length *L*, an image of the small illuminated patch appears as a shadow in the diffraction plane, magnified by a large factor *M*. Including spherical aberration in the analysis, Cowley showed that the magnification is modified to

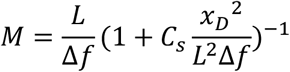

Thus, geometrical distortion starts to affect the shadow montage if *C*_*s*_*α*^2^/*Δf* is larger than a certain fraction (the reciprocal of the upsampling ratio defined further below). Typically, a 10% nonlinearity in magnification allows minimum defocus as summarized in table 1 (for a large spherical aberration, Cs=2.0 mm). In addition, the radius of the illumination cone at a certain defocus larger than *λ*/*α*^2^ is *αΔf*. The step size in real space, *Δx*_*S*_, should be smaller than this in order to ensure an overlap, which imposes an additional lower bound on the defocus, as shown in Table 1.

If we consider only the defocus aberration 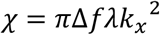 and a weak phase object approximation, where *k*_*x*_ is the reciprocal coordinate of the sample, then

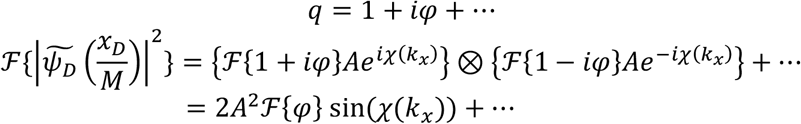

Thus, the contrast transfer function (CTF) is sin(𝒳) as for BF TEM (Fultz and Howe, 2002). For thicker samples, a similar analysis is based on a first order perturbation of the object function with respect to the average property. The phase contrast is dominant if the contrast inverts between overfocus and underfocus, whereas for still thicker samples the amplitude contrast dominates with a cosine transfer function.

It follows that the CTF flips sign first at 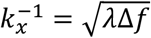 for phase contrast. (Using 200 kV and 1 μm defocus this figure corresponds to a spatial period of 1.6 nm at the sample plane.) Contrast will invert at certain higher spatial frequencies, similar to wide-field TEM. Mitigation for the distortion in representation of densities is achieved by choosing the smallest defocus that complies with Table 1, or by CTF correction.

### The shadow montage

The probe crossover during the scan should be consistently either above or below the sample, i.e., overfocus or underfocus. The degree of defocus should be larger in magnitude than both the depth of field unit *λ*/*α*^2^ and the spherical aberration length *C*_*s*_*α*^2^. Under this condition, undistorted shadow images are collected within the BF illumination disc. A shadow montage is simply the assembly of many image patches with an overlap consistent with the defocus, taking care that the summed intensity at each pixel should be normalized by the number of overlapping contributions. Importantly, different overlaps can be considered in parallel in order to address different values of defocus, effectively reconstructing a shadow volume based on different layers in the sample.

How can we predict the synchronization step corresponding to isolation of a layer at a certain defocus? According to the theoretical magnification, for every step of the probe by a distance *Δx*_*S*_, the shadow image at the detector plane should shift by a distance 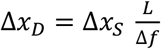.

The shadow image is inverted in × and y in the case of negative *Δf* (underfocus, meaning the probe cross-over is past the sampled layer). The number of pixels in the detector image to be shifted in each step is *Δx*_*D*_ times the density of pixels at the diffraction plane. If we count *N*_*BF*_ pixels as the diameter of the BF disc and the angular spread is 2*α*, then the spatial density of the pixels is *N*_*BF*_ /2*αL*. The result for the synchronization step (in units of pixels) is 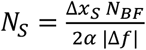.

The synchronization step *N*_*S*_ is also the upscaling factor of the rendered shadow montage, i.e. the ratio between reconstructed image and the scan sizes. Obviously *N*_*S*_ must be larger than one, and preferably 4 or larger, which imposes an upper bound for defocus that depends on the density of the diffraction space sampling. In addition, the choice of convergence angle and defocus should be bound by two limitations on resolution

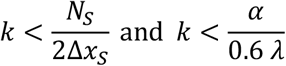

The first inequality states that resolution depends on the diffraction-pixel density of the detector. The second states that resolution of the up-sampled montage depends on the optical limit in the conventional way.

From the perspective of Figure 3 the shadow image is similar to a projection from a point source at the probe location. It is thus simple to understand how each layer contributes to the projection according to a different magnification determined by the distance from the point. This picture involves an implicit presumption that the exit-plane wave function is the product of a transmission function and the probe function, and therefore it is expected to break down under conditions of strong multiple scattering. Nonetheless, the geometric simplicity of the shadow concept suggests that the main effect will be a simple loss of contrast at the high resolutions rather than introduction of artefactual details. Synchronization between shadow images and the real space scan depends inversely on the defocus. This reflects the depth dependence of the magnification, but equivalently provides a route to optical sectioning. Given the presumption that a section is generated for each integer of *N*_*S*_, a depth resolution of *Δf*/*N*_*S*_ is expected, which is up to *N*_*S*_ times denser than the depth of field (see Table 1). The actual depth resolution may be further limited by the projection optics, detector resolution, and sufficient shadow overlap for decoupling between the sections. For a thin sample (order of *λ*/*α*^2^ or less), a single synchronization will suffice to provide an upsampling ratio relevant to the entire specimen thickness. For a thick sample and higher convergence, on the other hand, the synchronization, upsampling, and the defocus are coupled. This recalls the depth resolution enabled by multi-slice ptychography, according to a simplified explanation of the back-propagation in 3PIE (Maiden et al., 2012). A series of shadow montage reconstructions extending from one surface of the sample to the other could be called a shadow volume.

**Figure 3.**
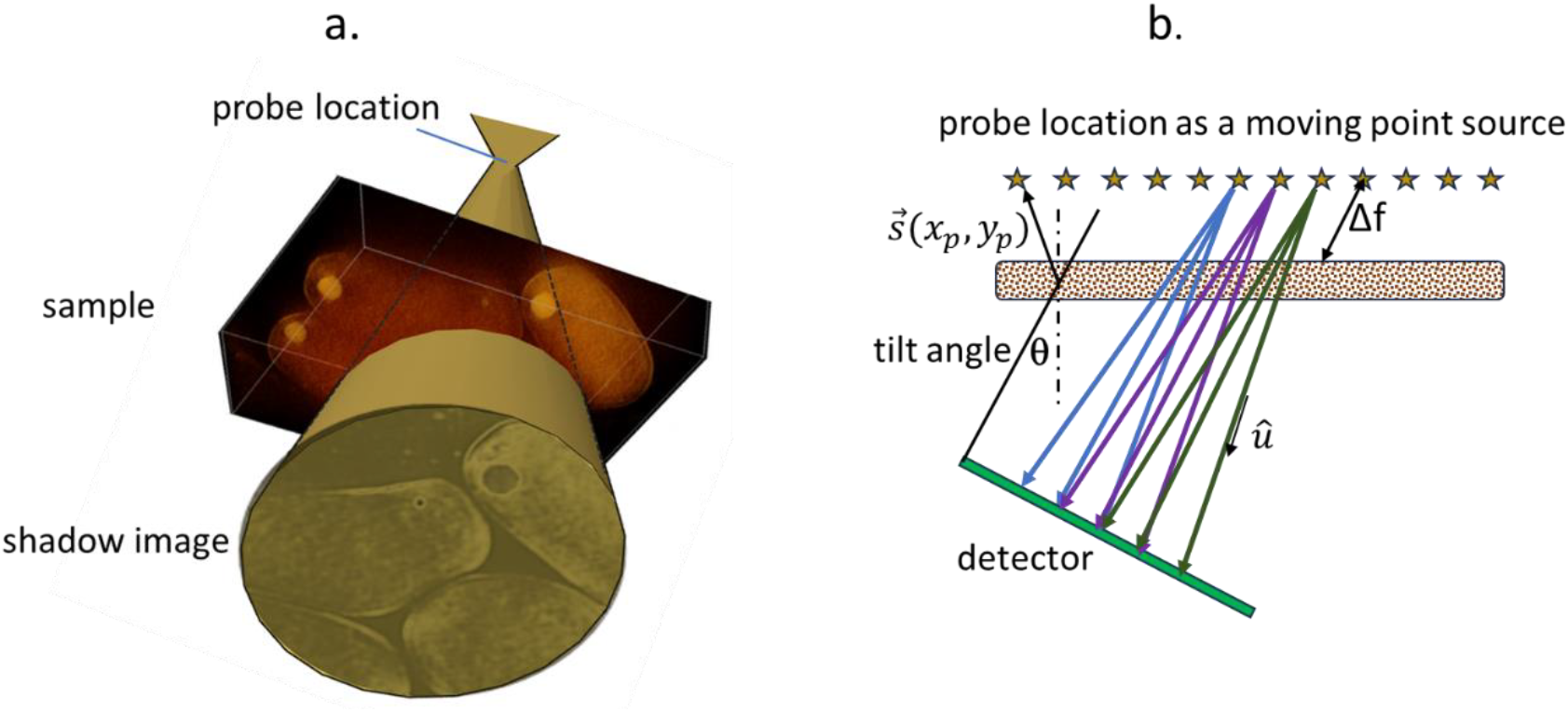
**(a)** The shadow image formed by a defocused probe is similar to a cone-beam projection. **(b)** The scanning probe can be considered a point source of illumination 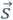 that follows a trajectory parallel to the sample. The different magnification of each layer in the sample is explained easily by the ray optics of the cone beam.

Our shadow montage may also be considered as a special case of a cone-beam back-projection reconstruction. Following (Bronnikov, 2000) we define a ray transform as a path integral from the source 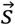 toward the sample along a unit vector 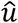

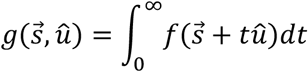

The back-projection reconstruction is the integration of ray transforms over a trajectory of the point source. A circular trajectory as shown in (Bronnikov, 2000) is popular, but the case of a planar scanning probe is valid as well. Reconstruction of the sample property *f*(*r*) according to the scheme shown in Figure 3b is achieved by

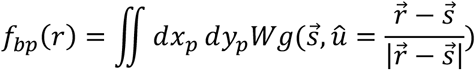

where 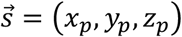 are the probe coordinates and W is weighting factor that in our case is W=1. In general, the weighting factor is tuned to allow a relation between the back-projection and the actual field according to convolution with a shift invariant point spread function, i.e.

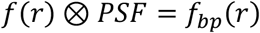

We note that for a particular layer of the sample and a particular position of the probe, the vector 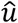 is mapped to a position of a pixel in the shadow image and the pixel intensity provides a value of the ray transform *g*. Thus, in the case of the shadow montage the integration required for the cone-beam back-projection is fulfilled by the overlap of many shadow images. As a special case of the cone beam, the shadow montage can in practice be implemented more efficiently.

The shadow volume or cone-beam reconstruction methods may also be combined with conventional tilt-series tomography. In principle, it should be possible to achieve a volume reconstruction uninterrupted by discrete angular sampling, similar to the implementation of a focal series acquisition at every mechanical tilt (Hovden et al., 2014). The major challenge in tomography is to find a set of transformations such that the raw data are described well with respect to rotation of a rigid body around a unique tilt axis. Normally this involves 2D alignment of the original projection images. This is justified for parallel projections (common practice in conventional TEM), or for the single-slice synchronization described above. For thicker specimens, determination of transformation for the tilt series alignment may be accomplished either using a reconstruction of the center slice or by alignment in 3D of the shadow volumes or cone-beam back-projections.

Finally, a 3D deconvolution on the back-projection result is recommended (Seifer, 2023). The 3D deconvolution aims at recovering *f*(*r*) from *f*_*bp*_(*r*), which in practice suffers from artifacts that originate in the limited and discrete angular tilt sampling. In practice, deconvolution suppresses the “fanout” artifacts recognized as projections of intensity into neighboring planes, and thereby improves the contrast in the (x,y) planes by removal of such cross-talk noise.

## Experimental

### Setup of Tecnai T20-F with ARINA detector and cryo-tomography

Data were collected using a Tecnai T20-F (FEI, Inc) transmission electron microscope operated at 200 keV in a 4D-STEM tomography mode with SerialEM (Mastronarde, 2003) and custom SavvyScan system as described previously (Seifer et al., 2021, 2024b; Seifer and Elbaum, 2023). The BF disk was directed to the center of the ARINA pixelated detector (DECTRIS SA, Baden-Daettwil, Switzerland) with a diffraction magnification (camera length *L*) chosen to fill a diameter typically covering 1/6 to 1/2 of the camera sensor, which was operated in binned 96×96 pixel mode to enable fast readout. In parallel, a dark-field signal was acquired on a HAADF detector (Fischione model 3000) mounted above such that its shadow would not interfere with the 4D STEM data. Each scan of 1024 × 1024 probe locations was recorded with spacing of 2.3 nm or 1.5 nm. The defocus was chosen to obtain a figure of synchronization *N*_*S*_ (i.e., upsampling ratio) typically between 3 and 10. Dynamic focus was implemented to maintain a constant defocus even as the sample plane tilts to high angle. Tilt series were acquired with steps of 3° typically between -60° and 60° in a dose symmetric manner (from lower to higher angles grouping 3 angles in each side in sequence). The scan pixel dwell time was 16 μs and the fluence was 2-3 electrons per Å^2^ per tilt view, so as to maintain a limit of approximately 100 *e*^−^/Å^2^ for an entire tilt series.

### Setup of Titan Themis with ELA detector

Zero-loss filtered 4D-STEM datasets were also acquired using a Titan Themis transmission electron microscope (Thermo Fisher Scientific Microscopy Solutions, Hillsboro, USA) equipped with a CEFID energy filter (CEOS GmbH, Heidelberg, Germany) and an ELA pixelated detector (DECTRIS SA, Baden-Daettwil, Switzerland). The microscope was operated at 200 keV and a semi-convergence angle of 8.6 mrad. Real-space images of 200×200 pixels with 4 nm spacing were collected, with diffraction magnification (camera length) chosen to project a BF disc of 196 pixels diameter on the camera image of 256×256 pixels at 2× binning. The physical size of the illumination cone on the detector is 29.4 mm, meaning that the effective camera length is L=1740 mm (which may differ from the nominal camera length).

### Biological Specimens

Tissue culture cells were prepared as previously reported (Kirchweger et al., 2025) by growth directly on gold Quantifoil grids (R2/2). The grids were glow discharged and coated with human fibronectin (FAL356008, Corning; 50 μg μl−1) in phosphate buffered saline without Mg2+ or Ca2+ for ∼1 h. Subsequently, 2 ml of growth medium were added, and the cells were seeded on the grids and returned to the incubator for growth overnight. Cells were vitrified by plunging to liquid ethane using a Leica EM GP plunger (Leica Microsystems, Austria) with the chamber held at 37°C and 95% humidity. 15 nm gold fiducial markers were added, and the grids were blotted from the back side for 5 s immediately before plunging.

### Data Processing

The data processing code for shadow montage assembly is written in Matlab and is publicly available in a GitHub repository: [https://github.com/Pr4Et/Supplementaries/tree/main/Shadow_Montage].

CTF correction may be applied optionally on the assembled shadow montage. (Applying a CTF correction on individual shadow patches is not recommended due to artifacts generated in the overlap process.) Processing time of the shadow montage on a Xeon workstation with 256 GB memory was 3 minutes per tilt view. 3D cone-beam back-projection routines from the Astra toolbox (van Aarle et al., 2015) were incorporated for comparison with the shadow montage. Spectral power was used as an indicator of image sharpness for tuning of the synchronization step parameter, evaluated as the mean spectral power in a Fourier ring covering between 1/8 and 1/100 of the image scale.

Tilt series alignment was performed with standard software, AreTomo (Zheng et al., 2022) or IMOD (Mastronarde and Held, 2017), using the shadow montage corresponding to the center of the specimen volume, or to the shadow volume. For a thin specimen (relative to the depth of field divided by the upsampling) the conventional back-projection or arithmetic methods (SART, SIRT) are justified. For a thick specimen, the entire depth cannot be rendered in focus without considering the depth-dependent magnification. This may be approached as a sum of the volumes generated by shadow volume or cone beam back-projection normal to the direction of the tilt plane (Radermacher, 2006). The alignment and back-projection steps may be seen as a synchronization and summation following the SSD paradigm proposed in the introduction.

## Results

### Shadow montage

Figure 4 compares a single projection image acquired with the Tecnai/ARINA setup to that acquired simultaneously on the HAADF detector. The specimen is a cultured tissue cell (U2OS) grown directly on the gold EM grid and recorded from an area of cytoplasm containing large vesicles. The following conditions were used: pixel sampling 1024×1024 @ 2.4 nm/pixel, semi-convergence angle α=4 mrad, diffraction image diameter *N*_*BF*_=40 pixels. Shadow montage with synchronization steps of *N*_*S*_ ≥ 6 appeared sharp, which indicates a defocus of 1.6 μm and therefore a patch radius of 6.4 nm. Panels a and b show the same fields of view for the HAADF and re-sized (downsampled) shadow montage, which is noticeably sharper. While the HAADF reflects probe resolution out of focus, the shadow montage is limited by the fundamental resolution of 0.6*λ*/*α*, the upsampling factor, and the noise. At electron dose of 3 *e*^−^/Å^2^ the up-sampled 6144×6144 @0.4nm/pix image is poor in contrast and requires downsampling to 2048×2048 to observe the muti vascular body seen near the center of Figure 4b. Downsampling a noisy image effectively averages neighboring pixels, reducing random fluctuations. According to the Central Limit Theorem, the standard deviation of uncorrelated noise decreases proportionally to the square root of the number of pixels averaged. As a result, the signal-to-noise ratio improves, and image contrast becomes more pronounced. The degree of downsampling is chosen as a trade-off between noise suppression and resolution loss, aiming to retain as much structural detail as possible.

**Figure 4.**
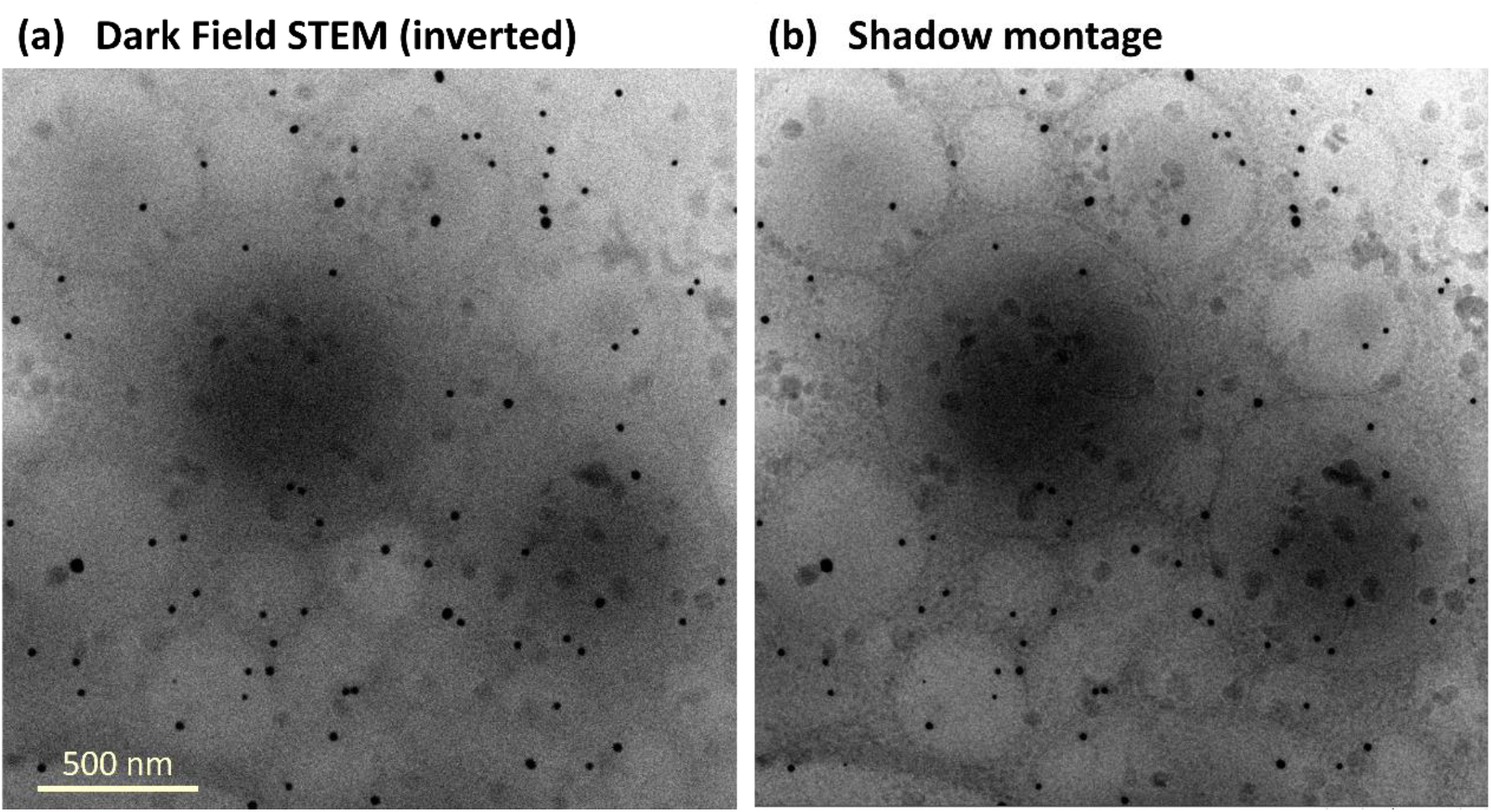
Comparison of **(a)** basic STEM using HAADF detector and **(b)** shadow montage by 4D-STEM using ARINA pixelated detector for a single tilt view. Also available in Supplementary Video 6.

Information acquired from a shadow image can be decomposed into the sum of BF intensity and the variations within the shadow normalized by the sum. The latter component characterizes the incremental information gained through the method, and can be isolated by normalization of the shadow patch image by the sum prior to assembly of the montage. Figure 5 displays an area of cell cytoplasm (G-8 cultured skeletal muscle cells). Panel a is the standard shadow montage and panel b shows the isolated variations. The illumination semi-convergence was α=0.8 mrad with *N*_*BF*_=47 and sampling step Δ*x*_*s*_=2.4 nm/pix. A focal series was acquired; for the displayed image acquired with underfocus of approximately -14 µm we predict a synchronization step *N*_*s*_=5. By scanning trial synchronization steps, Figure 5c, we find a maximum in the spectral power at *N*_*s*_ =4, and also that the precise value is rather forgiving at least for such small α. Comparison with a simple BF image acquired in focus, Figure 5d, shows that visibility of membranes is greatly enhanced in the shadow montage, and that the normalization behaves effectively as a background subtraction. Also evident in the images is contrast from surface contamination, which is equally enhanced by the normalization processing. Figure 5e shows the shadow montage for scan at overfocus of +14 µm, demonstrating inversion of the contrast from the shadow variations, whereas the background from the BF sum is not inverted (the large organelle volumes remain dark). Finally, we compare in Figure 5f with the parallax-filtered integrated differential phase contrast (πDPC) method previously reported (Seifer et al., 2024b), which was generated from the underfocus dataset. The low spatial frequencies are clearly observed and also part of the membranes are resolved, despite the significant defocus. The πDPC resembles acquisition with a virtual 2X2 pixel detector, and despite its sensitivity to depth information it cannot achieve the high fidelity of the shadow montage in recovering the finest features.

**Figure 5.**
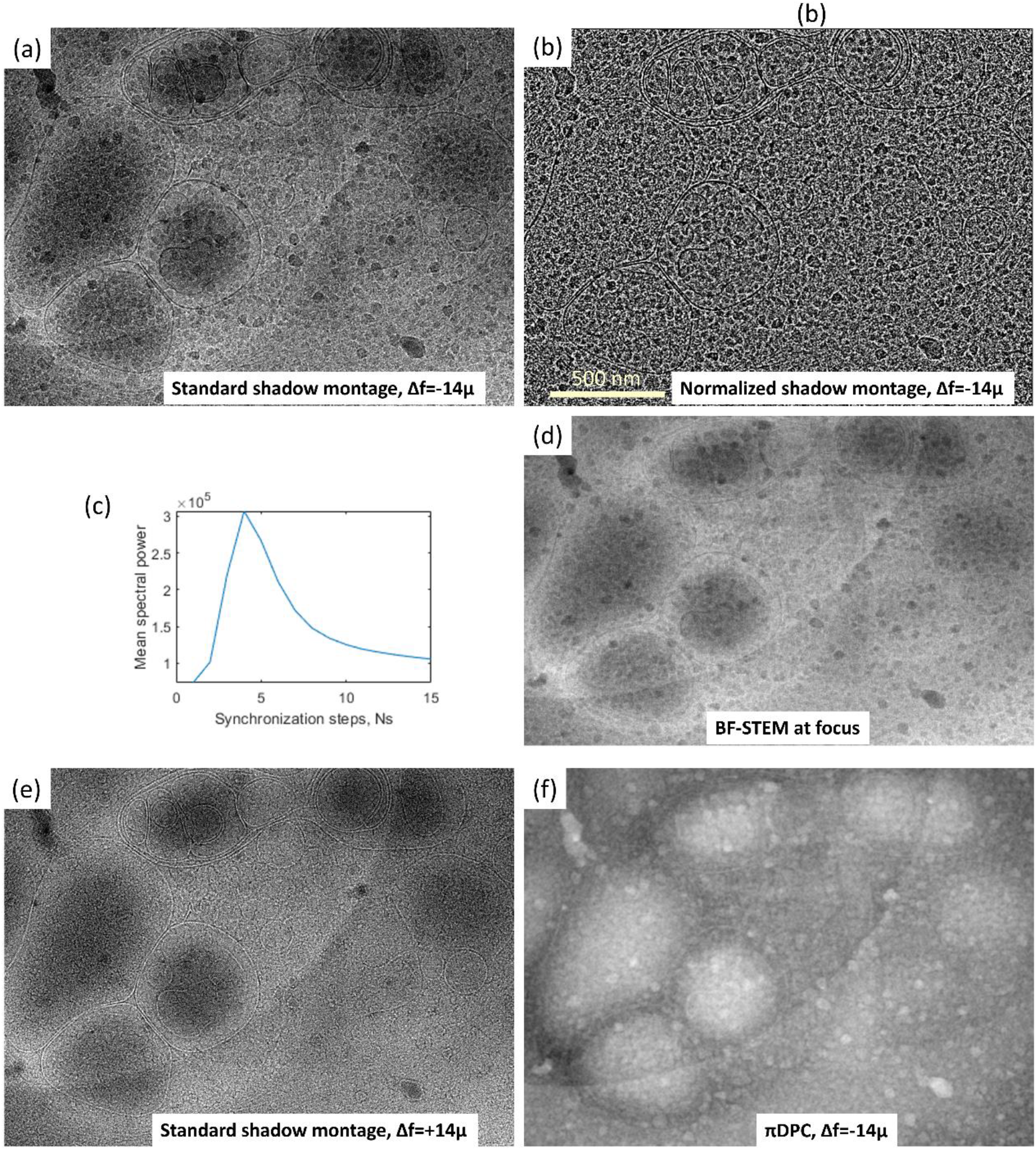
**(a)** A shadow montage acquired at 14 μm underfocus. **(b)** Patches normalized to a fixed BF electron count - leaving membrane and ice crystals visible owing to resolution of the pixelated detector. **(c)** Ns=4 was used to render the sharpest image according to the spectral power plot. **(d)** BF image of another dataset acquired at focus. **(e)** A shadow montage acquired at 14μm overfocus, flipped horizontally and vertically to match the image of panel a. The highest spatial frequencies are inverted, with membranes appearing bright as expected for a phase contrast transfer function. **(f)** The parallax-filtered integrated DPC of the underfocus dataset reveals long range order and part of the membranes.

Figure 6 compares shadow montages at synchronization steps *N*_*S*_ between 1 and 4 for a 4D-STEM dataset showing T4 bacteriophage analyzed previously with other contrast approaches (including πDPC and principal component analysis) (Seifer et al., 2024b). The projection dataset was recorded during a tilt series with semi-convergence α=0.8 mrad, a diffraction disc of diameter *N*_*BF*_=18, and real-space sampling 1024×1024 @ Δ*x*_*s*_=1.5 nm/pix. The limiting optical resolution is 0.6*λ*/*α*=1.9 nm, such that upsampling to a factor of 2 should reach the formal Nyquist criterion. Thus, the shadow montage images with upscaling *N*_*S*_ are resized to 2048 x2048 (upscaling by 2) to avoid false magnification. Nevertheless, there is a dramatic improvement in visible sharpness for synchronization steps *N*_*S*_ increasing from 1 to 3, and a marginal improvement for *N*_*S*_=4 that is visible only on close inspection. Our interpretation is that during recording the probe was unintentionally defocused by 4μm, matching synchronization of *N*_*S*_=4. The layers that synchronize with *N*_*S*_ =1,2,3 were approximately 17, 8.4, 5.6 μm away from the focal plane and thus appear blurred. The clarity of the shadow montage at the optimal *N*_*S*_ exceeds that of previous methods applied to the same dataset, which encourages our SSD strategy.

**Figure 6.**
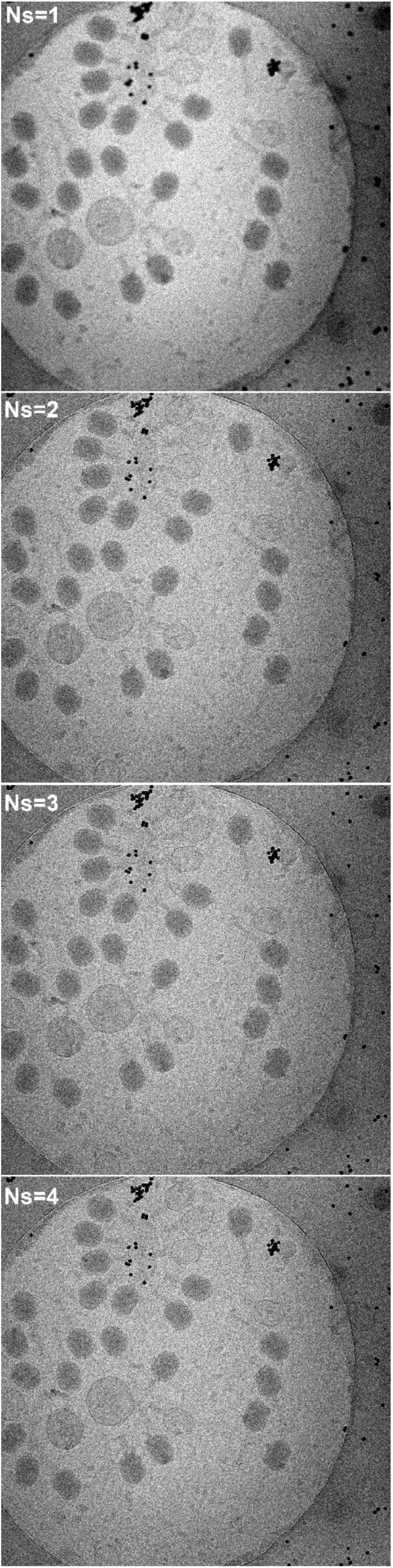
Increasing the synchronization step N_S_ up to 4 improves the sharpness of details in the shadow montage of a T4-bacteriophage sample.

### Depth resolution

Depth resolution in a convergent projection was demonstrated using a specimen of colloidal gold particles dispersed on a support film in the Titan Themis setup with ELA detector. As often happens, the film ruptured in places and folded back upon itself so that the gold particles were suspended in two distinct planes representing different depths. The aberration-corrected probe (α=8.6 mrad) was defocused by 2.8 μm and 2.1 μm with respect to the two carbon layers. The synchronization steps to make the two layers appear sharp were *N*_*s*_=16 and *N*_*s*_=22, in accordance with our analytical formula (see Figure 7a). A sweep through synchronization steps between 12.6 and 24.6 is found in Supplementary Video 3, which demonstrates the resemblance to a virtual focus control post-acquisition.

**Figure 7.**
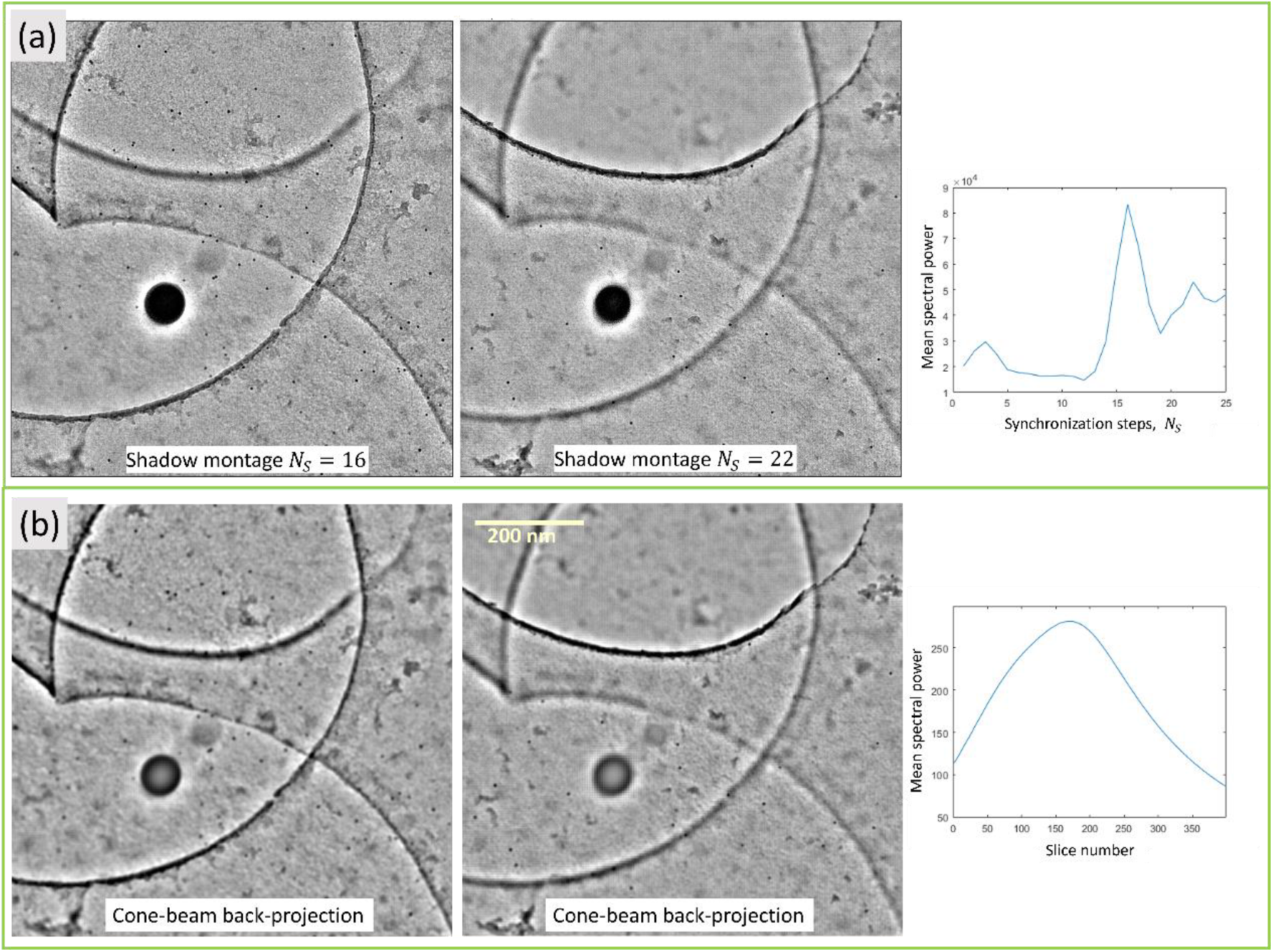
Two carbon layers covered with 10u gold particles acquired in a single 4D-STEM scan. **(a)** Separable by the shadow montage with different synchronization steps. The effect is fully appreciable by watching a sweep through more layers shown in Supplementary Video 3. **(b)** The layers are also separable by a cone-beam back-projection generated using Astra toolbox. Note the different gold particles that appear sharply in each layer.

Supplementary Video 4 shows a similar scan recorded without the energy filter. The two synchronization steps are also recognized as peaks in the mean spectral power of the image. The overall count of electrons is 4 times higher in the non-filtered case in accordance with the difference between elastic and inelastic scattering electron count in carbon up to the illumination angle of 8.6 mrad (Seifer et al., 2024a). At the level of a visual inspection the results appear identical, with or without the energy filter. However, the energy-filtered case demonstrates better separation between the layers comparing spectral power plots in the figures.

As explained above, the shadow montage is theoretically equivalent to cone-beam reconstruction with a grid of source points on a plane. Figure 7b displays a 3D cone-beam back-projection reconstruction, prepared using a module in Astra toolbox. This module excels in solving the most general case of cone-beam reconstruction. It is not optimal, nor strictly designed, for the case of a planar grid with millions of projections. Nevertheless, the code succeeds in generating a volume of 400 layers with lateral resolution of 1024x1024 (limited by the GPU memory). The nanoparticles are clearly visible in the two layouts, although the spectral plot does not recover two distinct peaks because of the lower resolution. Being a more specific task, the shadow montage outperforms the cone beam reconstruction in execution time and memory usage. Yet, essentially, the operations are very similar.

### A difference between BF TEM and shadow montage

The relative insensitivity to inelastic scattering is often quoted as an advantage of STEM imaging with respect to conventional wide-field TEM. In the latter case, the objective lens chromatic aberration introduces a defocus spread; this is normally addressed by zero-loss energy filtering, which results in a loss of signal intensity. In STEM, the resolution-limiting probe focus occurs prior to interaction with the specimen, and the effect of inelastic scattering appears in the angular distribution at the diffraction plane (Seifer et al., 2024a). To understand this more deeply, we collected at fixed defocus atomic resolution shadow images series from a thin sample of tungsten diselenide, while detuning the acceleration voltage on one hand, or the energy filter liner tube voltage on the other (see Supplementary Video 5). In both cases the detuning was over a range of 0 - 40 V, so as to cover the plasmon peak seen prominently in biological cryo-EM. The change in illumination wavelength results in a magnification change that is most evident in the FFT. The change in liner tube voltage, on the other hand, affected the atomic visibility due to actual energy losses in the specimen, but the FFT pattern was unaffected. These results indicate that the shadow image maintains the STEM-like insensitivity to electron energy losses occurring after the probe focus. At the same time, we conclude that tuning of the synchronization steps cannot filter for energy losses in the specimen.

### Shadow Montage Tomography

Upsampling of the 2D projection images may be exploited to improve the resolution of tomographic reconstruction. Figure 8 shows the reconstruction of the bacteriophage tiltseries dataset in tilt views between -60° and 60° at steps of 3° (of which the center view is shown in Figure 6), synchronized with *N*_S_=4 and displayed in orthogonal sections covering a selected sub-volume. For comparison, we display the same volume processed by the previously-developed parallax-filtered integrated differential phase contrast (πDPC), which does not benefit from the upsampling.

**Figure 8.**
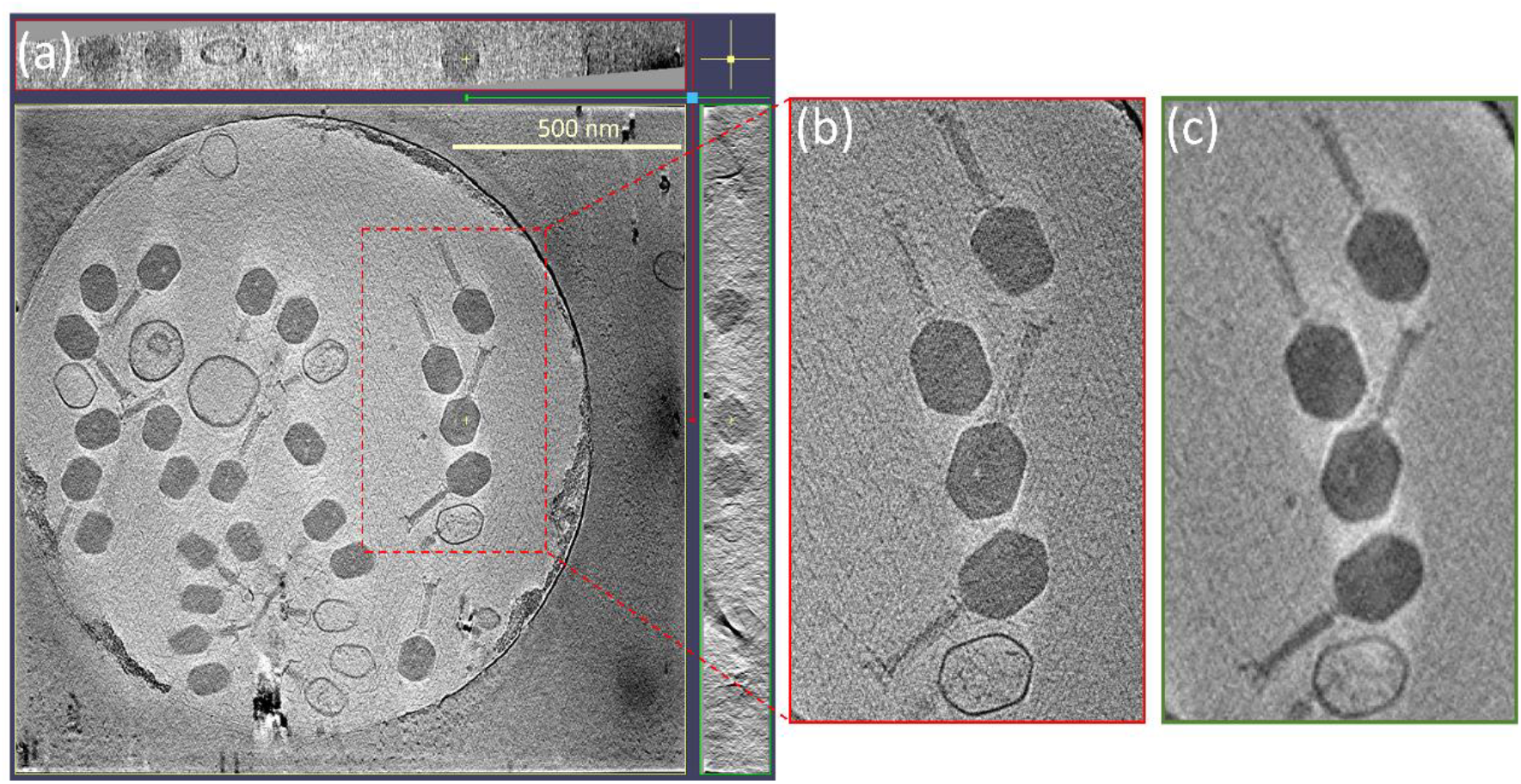
**(a)** 3D reconstruction from a tilt series of shadow montages such as shown in Fig.7 for N_S_=4 with final rendering to twice the scanning probe resolution. The different phases of the T4-bacteripogaes are clearly observed. Also available in Supplementary Video 7. **(b)** Close up on one area. **(c)** A similar reconstruction from πDPC projection images, at the scan resolution.

The next conceptual advance is to combine the depth-dependent shadow volume reconstruction *per tilt* with the conventional tilt tomography scheme covering a wide range of angles. This is demonstrated using a tilt series acquired from an area of rough endoplasmic reticulum in the cytoplasm of a MEK (mouse embryonic kidney) cell cultured on the EM grid. The shadow volumes were constructed from automatically determined *N*_*s*_ steps per tilt view based on defocus estimation and thickness, which were further processed by interpolation and geometric correction to form dense layers. AreTomo2 (Zheng et al., 2022) was used with the center slices for alignment. Figure 9a shows the default reconstruction by the SART method, which presumes parallel projections. Figure 9b shows the reconstruction based on back-projection from the series of 3D shadow volumes. Figure 9c displays the back-projection after 3D deconvolution (Seifer, 2023).

**Figure 9.**
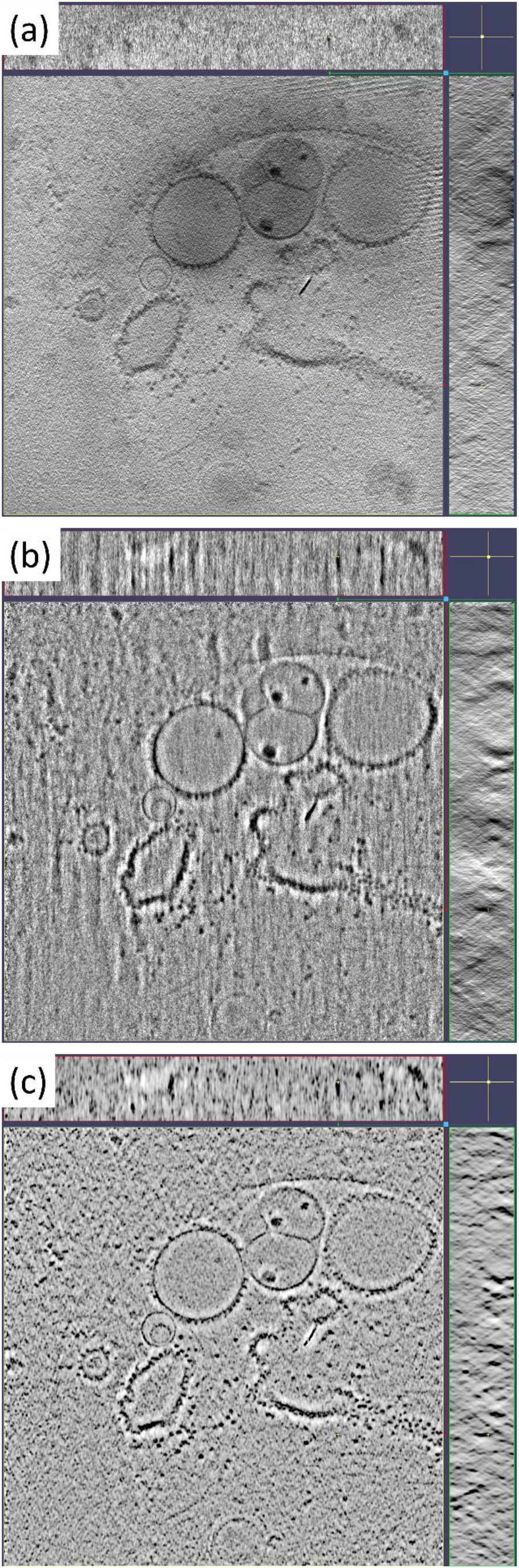
Tomograms based on shadow montage images of MEK cell sample. **(a)** Standard reconstruction using AreTomo2 on generated tilt series from shadow montage. **(b)** Volume (multi-slice) shadow montage followed by back-projection. **(c)** 3D deconvolution of the back-projection. (Detailed reconstructions appear in Supplementary Videos 8-10.)

## Conclusions

We have demonstrated the reconstruction of a bright-field image from far-field diffraction data recorded with a defocused scanning probe. Contrast transfer is similar to that of wide-field TEM, and the montage generation may be considered as a simplified implementation of the tcBF method on one hand, or of the cone-beam back-projection on the other. There is also an obvious analogy to the spot scanning method used in film-based TEM (Downing, 1991), but the shadow image maintains the advantage of insensitivity to energy loss in the specimen.

In comparison with classic STEM methods that operate in focus, and in common with the tcBF, the shadow montage offers a means to acquire a large field of view at the limiting optical resolution with a relatively small number of probe positions and with greater tolerance on the focus tuning. The image processing is straightforward and direct, in the sense of avoiding the need for iterative algorithms whose convergence may be very problematic for sparse data. Only the defocus must be determined in order to set the synchronization parameter *N*_*s*_. This may be done by comparison of image shifts as described for tcBF or πDPC, or empirically by scanning a range of trial values and evaluating the spectral power curves. Looking forward, if higher order aberrations are known, or can be determined from the data, it should be possible to correct them by affine transformation of the shadow patch prior to stitching; defocus correction is only the simplest case because it presumes a uniform magnification independent of *k*. The output of the shadow montage processing may serve as well as an input for refinement by full iterative ptychography. The full power of the shadow montage should be revealed in application to tomography of moderately thick specimens. The possibility to refocus computationally to different depth, shown most vividly for the gold on carbon sample simply by choice of synchronization steps, offers a means to reach high lateral resolution without restriction of the parallel projection presumption that underlies the tomographic back-projection reconstruction. Essentially there is a depth-dependent magnification, which might be addressed in full by multi-slice ptychography, or alternatively by cone beam back-projection or shadow montage synchronization, both in combination with conventional tilt tomography. Here we have demonstrated the protocols and their potential using an intact cell specimen, without the milling or thinning required for conventional TEM. The ease of implementation of the shadow montage suggests it as a method of choice for 4D-STEM imaging and tomography, particularly for radiation sensitive objects. We also expect that other users of cone-beam reconstructions may find the shadow montage a useful processing approach.

## Data Availability

All the electron microscope datasets are publicly available on Zenodo (Seifer, 2024; Seifer et al., 2025a, 2025b).

## Acknowledgement

Arina Dalaloyan and Peter Kirchweger contributed in the preparation of samples. The authors acknowledge support from the Irving and Cherna Moskowitz Center for Bio and Nanobio Imaging, and from the European Union (ERC-Adv CryoSTEM, 101055413; Views and opinions expressed are however those of the authors only and do not necessarily reflect those of the European Union or the European Research Council. Neither the European Union nor the granting authority can be held responsible for them.) ME is incumbent of the Sam and Ayala Zacks Professorial Chair in Chemistry. This research is made possible in part by the historical generosity of the Harold Perlman family.

## Notes

### Competing Interest Statement

The authors have declared no competing interest.

