## Supplementary material for "Shadow montage and cone-beam reconstruction in 4D-STEM tomography": Figure S1

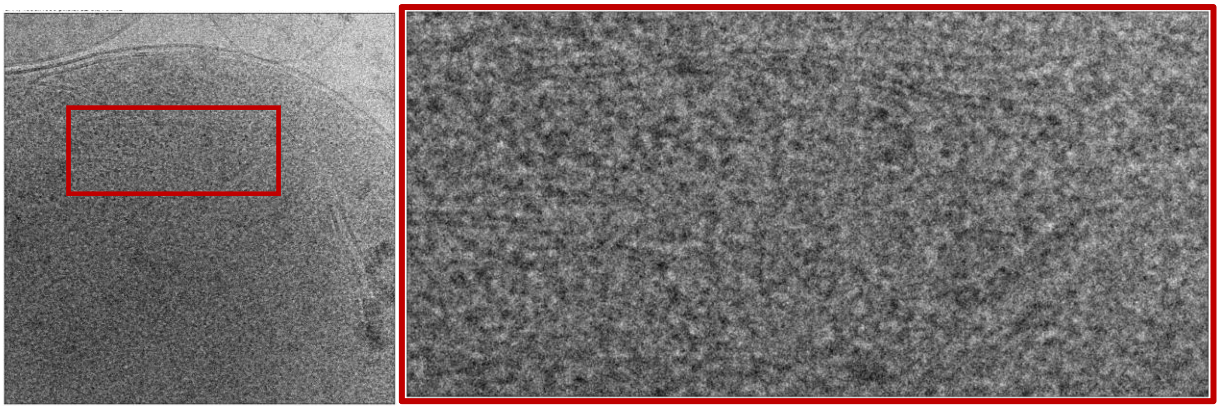

*Figure S1. Using shadow montage method to regenerate the tc-BF result presented in Figure 3a,c of Yu et al [https://www.biorxiv.org/content/10.1101/2024.04.22.590491v1]. Processed from dataset 42_scan_x256_y256 in https://zenodo.org/records/10825339 using synchronization steps 14 and 15 pixels, averaged.*
